## Supplementary figures and images for "Enhanced virulence and stress tolerance are signatures of epidemiologically successful *Shigella sonnei*"

### Supplementary figures

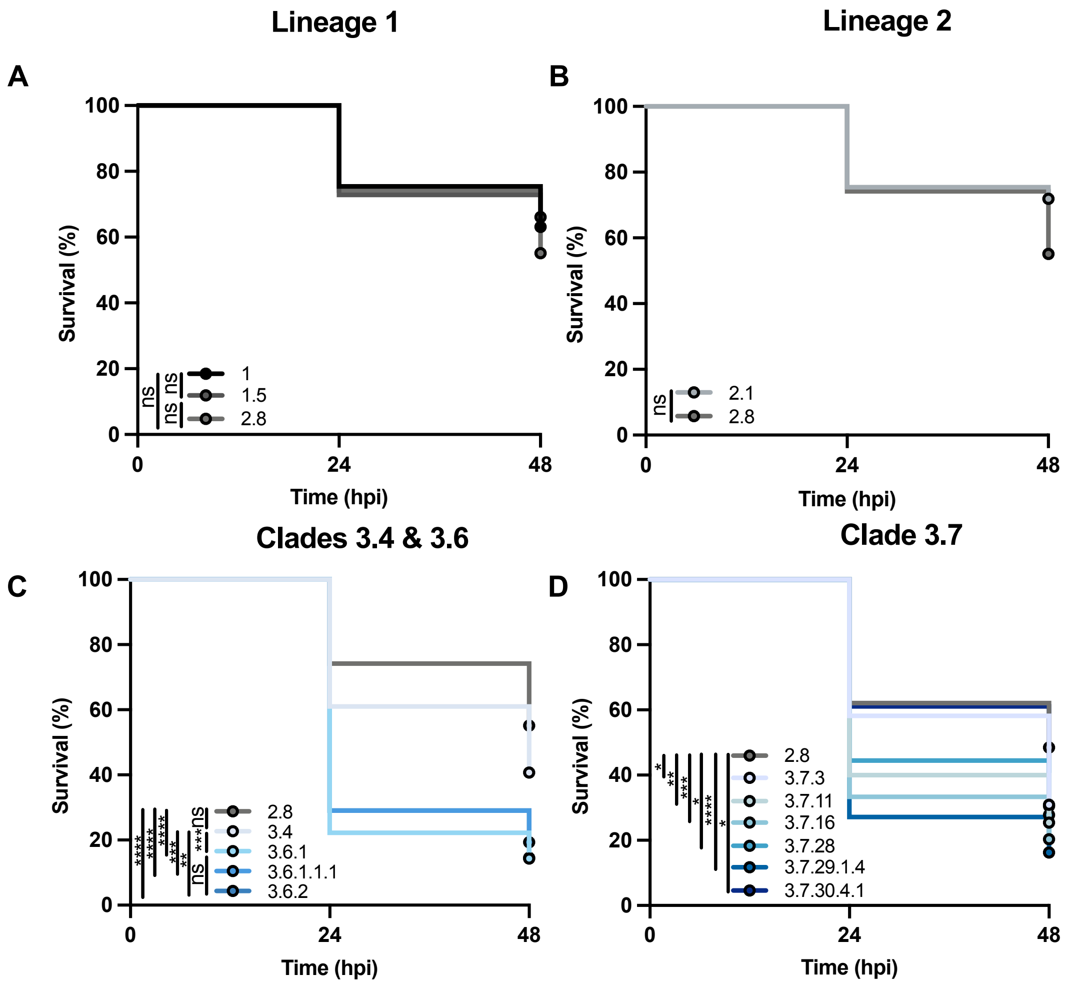

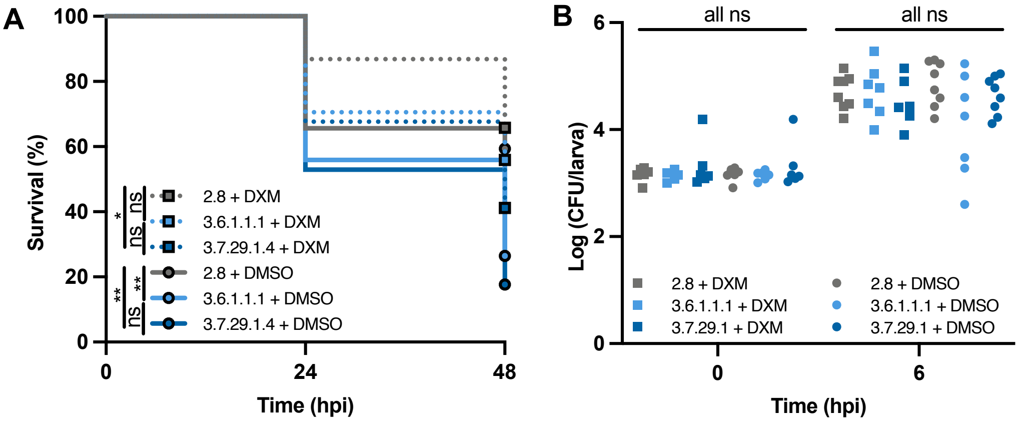

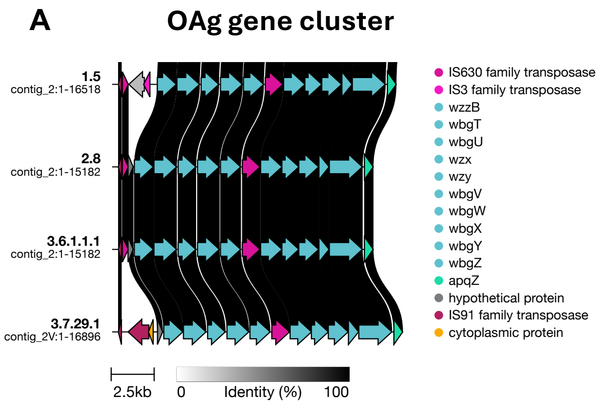

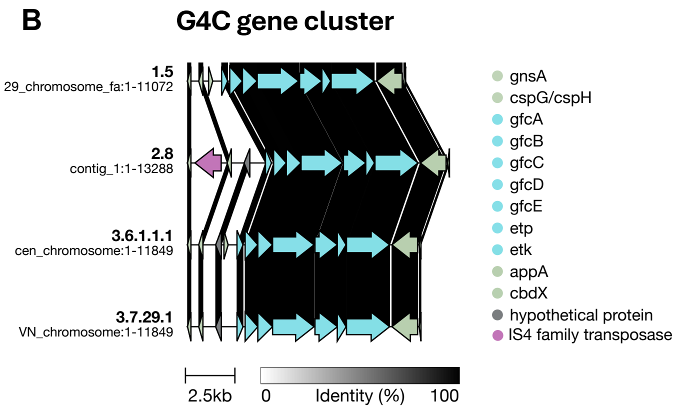

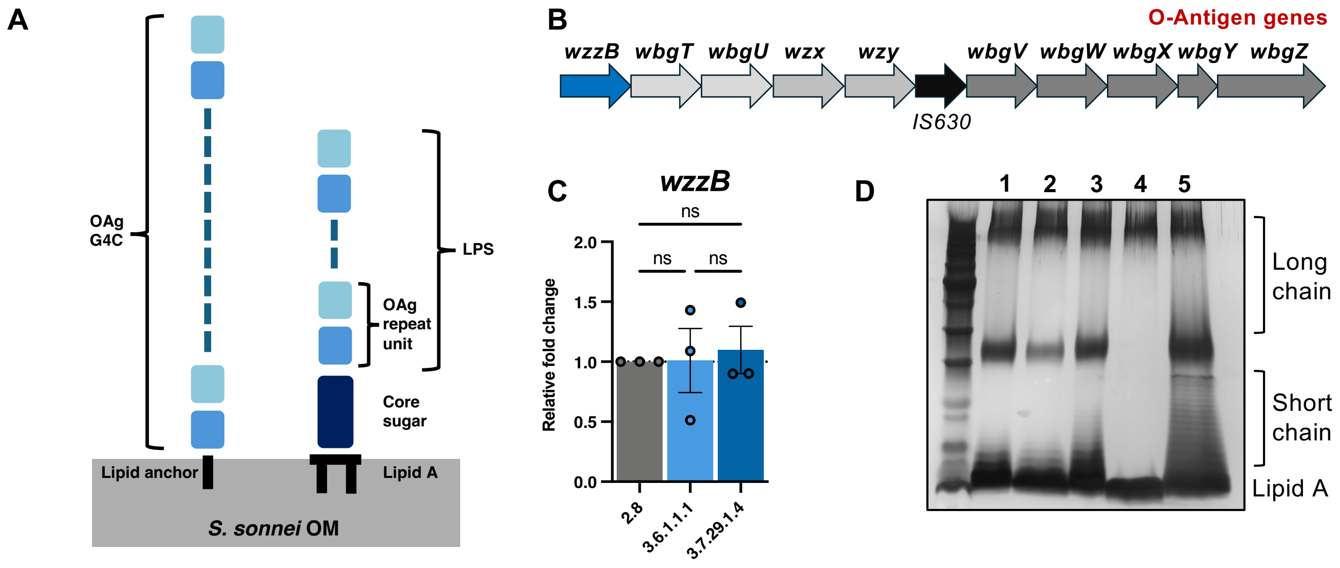

#### Figure S5

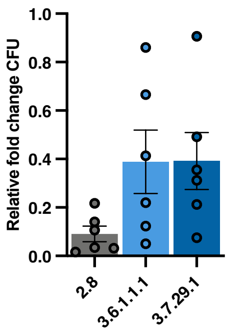
